# CRISPR-Cas9 based Mutagenesis in the Entomopathogenic Nematode *Steinernema hermaphroditum* and the Maintenance of Mutant Lines

**DOI:** 10.1101/2025.10.14.682448

**Authors:** Sally W. Ireri, Mengyi Cao

**Affiliations:** Division of Biosphere Sciences and Engineering, Carnegie Science, Pasadena, CA, USA

**Author notes:** Email address of the corresponding author: Mengyi Cao. Email address of the co-author: Sally W. Ireri.

## Abstract

Entomopathogenic nematodes (EPNs) from the genus *Steinernema* and *Heterorhabditis* form mutualistic relationships with symbiotic bacteria from the genus *Xenorhabdus* and *Photorhabdus*, respectively. Together, these nematode-bacterium pairs infect and kill insect hosts—primarily larvae from the orders *Lepidoptera* and *Coleoptera*. This tripartite interaction provides a powerful model for investigating the molecular mechanisms underlying mutualism and parasitism. A key step toward this goal is the development of a genetically tractable EPN. While RNAi has been applied in some EPN species, stable, transgenerational genetic tools remain limited. Here, we establish a robust CRISPR-Cas9 system in the emerging model *Steinernema hermaphroditum*, a species that is easily cultivated in both *in vivo* and *in vitro* conditions and amenable to gonadal microinjection. Notably, its hermaphroditic reproduction simplifies the generation of genetically stable mutant lines. We present a detailed protocol for efficient, targeted gene knockout via microinjection in *S. hermaphroditum*. As a proof-of-concept, we knocked out a conserved homologue, *unc-22*, which causes a twitching phenotype. The CRISPR-Cas9 based genome editing in *S. hermaphroditum* has potential to be used to express transgene, or to be adapted to other EPN species that are applicable to benefit agriculture.

**SUMMARY:** This article demonstrates CRISPR-Cas9 mediated genome engineering in *Steinernema hermaphroditum*, an entomopathogenic (EPN: insect-parasitic) nematode and an emerging genetic model. The described technology is useful for creating mutants allowing for the elucidation of gene functions in the nematode biology that is relevant to mutualistic and parasitic symbiosis.

## INTRODUCTION

Entomopathogenic nematodes (EPNs) are insect-killing parasites which form species-specific mutualistic partnerships with their bacterial symbionts^1^. EPNs consist of two families: Steinermatidae and Heterorhabditidae which associate with Gram-negative bacteria of *Photorhabdus* and *Xenorhabdus* spp, respectively^2, 3^. During the free-living infective juvenile stage (IJ) of *Steinernema* nematodes, *Xenorhabdus* bacteria are housed within an intestinal pocket of the nematode known as the receptacle and upon penetration of the insect host into the hemocoel the bacteria are released, multiply and kill the insect by septicaemia^3, 4^. In this mutualistic relationship, the bacteria produce antimicrobial and insecticidal compounds, which protect against the host immune response and provide nutritional support to the nematode^5^. In return, the bacteria benefit from the protection provided by nematode host against environmental microbes and are facilitated in their dispersal to new insect prey^6, 7^.

Over the past few decades, the *Steinernema-Xenorhabdus* partnership has been established as a powerful experimental system to study various aspects of parasitic and mutualistic interactions, such as insect host-seeking behaviors^3^, bacterial colonization in the nematode host^8^, and adaptive behaviors of symbiotic bacteria that facilitates their transitions between nematode and insect host animals^9^. EPNs are also important species in soil sustainability and used as organic pest control agents to promote agricultural productivity^10, 11^.

Despite the rapid development of genetic tools in the symbiotic bacteria *Xenorhabdus* and *Photorhabdus*^12^, consistent genetic tools in EPN have been scarce. Gonadal microinjection has been successfully established in both *Heterorhabditis* and *Steinernema* nematodes^13, 14^. Previously, gonadal microinjection in *Heterorhabditis bacteriophora* has been used to consistently deliver double-stranded RNA for gene knock-down among the first generation of progeny^13, 15^. Recently, we developed CRISPR-Cas9 genome editing in *Steinernema*, delivered by gonadal microinjection, and created precise, stable, and heritable mutations^14^.

One *Steinernema* spp, *S. hermaphroditum*, has been demonstrated to be highly tractable and has great potential as an emerging genetic model organism^4, 14^. Originally isolated from the soil in Indonesia^16^ and re-isolated from India^17^, *S. hermaphroditum* is consistently hermaphroditic, with a small number of males in each generation — two features that facilitate the adaptation of genetic tools from the classical model nematode *C. elegans*^4^. In addition, the genome of *S. hermaphroditum* has been sequenced and assembled into chromosomes, simplifying the task of identifying potential Cas9 target sites, and designing primers ^18^.

During microinjection of *Steinernema hermaphroditum*, an injection mix containing appropriate reagents, such as dsRNAs, Cas9 protein, or guide RNAs, are delivered to the syncytial nematode gonad via a needle pulled from a quartz capillary. The appropriate volume and pressure of injection result in a visible ‘flow’ of liquid throughout the gonadal arm. Since these nematodes have a pair of syncytial gonads composed of germline nuclei that share the same cytosol^19^, this microinjection approach allows multiple germ cells to be accessible for editing, before cytokinesis occurs separating the germline nuclei into developing oocytes^20^. Although gonadal microinjection serves as a consistent technique for genetic modulation in several species of non-model nematodes^14, 21–25^, it is crucial to adapt the injection techniques to the appropriate gonadal structures, life stage, and physiological condition of each nematode species to ensure the success and high efficiency in delivery. Recovery techniques post-injection and maintenance of mutant lines are both important to ensure this technique is cost-efficient.

Here we demonstrate a CRISPR-Cas9 genome editing protocol in *Steinernema hermaphroditum*. We describe the gonadal microinjection in the young adult, a stage that is resilient to high-pressure microinjection and is receptive to gene editing^14^. In addition, we provide detailed methods describing the maintenance of heritable and stable mutant lines both via cryopreservation and propagation through insects.

## PROTOCOL

### 1. Nematode preparation prior to microinjection

#### 1.1. Prepare Nematode Growth Media (NGM) plates seeded with bacteria *X. griffiniae* and *C. aquaticus*

Grow *S. hermaphroditum* nematodes on their native symbiotic bacterium *Xenorhabdus griffiniae* before microinjection to maximize nematode growth and recover injected animals on bacterium *Comamonas aquaticus* to increase post-injection survival^14^.

1.1.1. Streak bacterium *X. griffiniae* on LB-pyruvate agar plates (2.5 g yeast extract, 5 g tryptone, 2.5 g sodium chloride, 0.5 g sodium pyruvate, 7.5 g agar in 500 mL water) and incubate overnight at 30 °C.

1.1.2. Streak bacterium *C. aquaticus* on LB agar plates and incubate overnight at 37 °C.

1.1.3. Pick a single colony of bacteria to inoculate dark LB liquid media (5 g yeast extract, 10 g tryptone, 5 g sodium chloride in 1 L of water). Grow bacterial culture overnight (16 hours maximum) with aeration (*X. griffiniae* at 30 °C and *C. aquaticus* at 37 °C).

1.1.4. Prepare NGM agar: (3 g NaCl, 2.5 g peptone and 20 g agar added to 975 mL water, with 1 mL cholesterol, 1 mL 1 M CaCl2, 1 mL 1 M MgSO4, 25 mL 1 M KPO4 added after autoclaving) 10 mL per 60 mm x 15 mm Petri dishes.

1.1.5. Seed bacteria: add a few droplets of overnight bacterial culture onto each NGM agar plate, gently shake to spread the bacteria droplets into a patch.

1.1.6. Incubate bacteria-seeded NGM plates at room temperature to allow agar to dry and bacteria to grow.

NOTE: *X. griffiniae* seeded plates should be used as soon as possible; *C. aquaticus* plates can be stored at room temperature for 1-2 weeks.

#### 1.2. Nematode staging

Pick the J3 stage of hermaphrodite the night before injection and young adults on the day of injection.

1.2.1 Pick or chunk *S. hermaphroditum* nematodes onto the prepared NGM plates seeded with *X. griffiniae*. Incubate at a temperature of 25°C -28 °C for optimal growth.

1.2.2 The night before microinjection, pick J3 stage of hermaphrodites onto NGM agar seeded with *X. griffiniae*. As shown in Fig 1A, J3 stage is estimated by the size of the nematode and the morphology of gonads and vulva.

**Figure 1:**
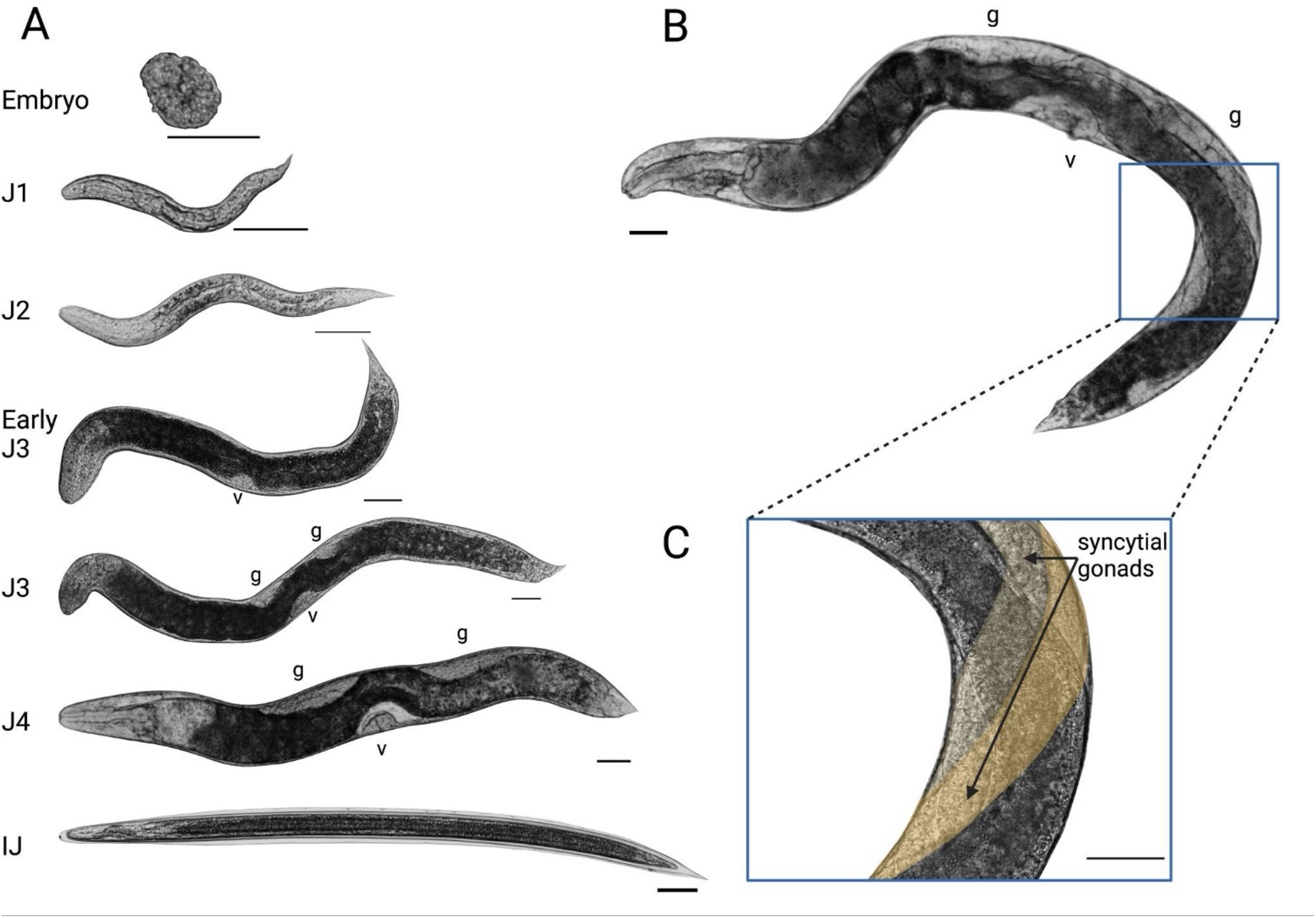
The development of *S. hermaphroditum* in a laboratory setting. (A): Estimated developmental stages of hermaphrodites grown on NGM agar feeding on its symbiotic bacterium *X. griffiniae*. Scale bar = 50 μM. v = vulva; g = gonad. (B): A young adult hermaphrodite at the appropriate developmental stage for gonadal microinjection. Blue box: location of gonadal injection. (C): The syncytial gonad of the hermaphrodite targeted for microinjection. The two gonads are colored in different shades of yellow to facilitate visualization.

1.2.3 On the day of microinjection, pick young adult hermaphrodites [Figure 1B].

### 2. CRISPR-Cas9 preparation for microinjection

#### 2.1. Selecting the CRISPR crRNAs and reagent storage

2.1.1 Extract the target gene sequence from the reference *Steinernema hermaphroditum* genome using BlastP, PRJNA982879.

2.1.2 Select an early coding exon sequence, if possible, preferably the first exon, and input the sequence into CRISPRScan using default settings ^26^ to generate a list of possible CRISPR RNAs (crRNAs).

2.1.3 Blast the top crRNAs against the *S. hermaphroditum* genome to ensure only unique crRNAs are chosen.

Optional: Confirm the sequence of the target region of the genome by PCR amplification and Sanger sequencing before crRNA design, to ensure efficient binding of crRNAs.

2.1.4 CRISPR RNAs (crRNAs) and universal trans-activating CRISPR RNAs (tracrRNAs) are stored at 100 μM each in duplex buffer at -80 °C.

2.1.5 Cas9 protein is stored at 10 mg/mL at -20 °C or -80 °C.

2.1.6 1 M KCl is stored at 4 °C.

#### 2.2. Making the 2% agarose pad for injection (adapted from Berkowitz, L.A *et al*.^27^)

2.2.1 Melt 2% agarose in water using a microwave oven.

2.2.2 Transfer 50 μL of melted agarose onto a cover glass (48 x 60 mm no.1). Immediately cover the agarose droplet with another cover glass.

2.2.3 Let the agarose dry at room temperature for 20 minutes. Remove the top cover glass. Label the front side of the agarose pad.

2.2.3.1 Optional: bake the agarose pad at 37 °C overnight to further dry the agarose pad.

2.2.4 2% agarose pads could be stored at room temperature for months to years.

#### 2.3. Needle preparation

2.3.1 Manufacture quartz needles using the following parameters on a laser filament puller. With the needle puller we use (see model in Materials list), the suggested program is listed below: Heat: 700 Filament: 4 Velocity: 60 Delay: 145 Pull: 175

2.3.2 Quartz needles (OD: 1.0mm, ID: 0.7mm) can be pulled immediately before injection, or ahead of time and stored in a pipette storage box.

2.3.3 Alternatively, we also have success injecting with borosilicate needles (OD: 1.2 mm, ID: 0.68 mm). Note that different sizes of needles will match different sizes of needle holder.

### 3. CRISPR-Cas9 microinjection and Recovery of the nematodes after microinjection

#### 3.1. Prepare the CRISPR-Cas9 microinjection mixture (adapted from Wang, H., *et al*.^28^)

3.1.1 Prepare fresh guide RNA duplexes by mixing the chosen crRNAs and universal tracrRNA in a 1:1 ratio, i.e., total crRNA to tracrRNA in a PCR tube:

|  |  |
| --- | --- |
| crRNA #1 (100 μM) | 1.5 μL |
| crRNA #2 (100 μM) | 1.5 μL |
| tracrRNA (100 μM) | 3.0 μL |

NOTE: If using co-CRISPR, crRNA #1 is targeting *unc-22* marker gene, while crRNA #2 can be replaced with another sequence targeting a second gene of interest.

3.1.2 Incubate the PCR tube in a thermocycler at 94°C for 2 min and then cool to room temperature.

3.1.3 Assemble the injection mix by combining the following, then incubate for 5 min at room temperature:

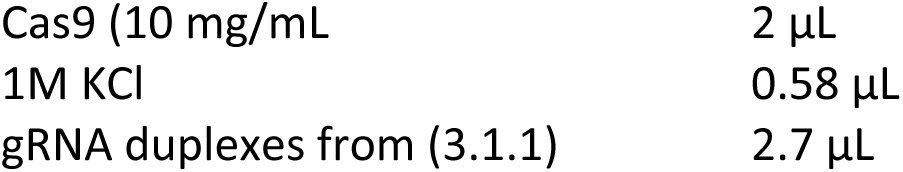

3.1.4 Mix three times by flicking the PCR tube followed by a brief spin (10-15 seconds) in a mini centrifuge to collect the contents of the tube. After the final mix, spin down for longer, for up to 1 minute.

#### 3.2. Load and break the needle

**3.2.1**. Load the needle with 1-2 μL of injection mixture using a microloader tip.

**3.2.2**. Assemble the needle to the pipette (needle) holder then break open the tip by dragging the needle across a piece of double-sided tape covered in halocarbon oil. Test the opening of the needle using the ‘CLEAN’ function of the microinjector.

#### 3.3. Microinjection into the syncytial gonads of young adult nematode

3.3.1 Transfer a small droplet of **Halocarbon oil** onto 2% agarose pad.

3.3.2 Under the dissecting stereoscope, use a thin (soft) platinum wire to pick up a young adult hermaphrodite nematode and wash it in an M9 buffer (per liter: 3 g KH2PO4, 11.3 g Na2HPO4, 5 g NaCl, with 1 ml 1 M MgSO4 added after autoclaving) to remove bacteria (see young adult nematode in Figure 1].

3.3.3 Transfer the washed nematode onto an agarose pad and place it in the droplet of halocarbon oil. Use a soft platinum wire to gently brush the nematode until the animal is immobilized.

3.3.4 Carefully place the agarose pad on the microinjection stage: the orientation of the gonadal syncytium should be pointing to the needle at about 45° angle. Start from 5x magnification, then switch to 20x and 40x magnification for the injections.

3.3.5 Ensure the needle penetrates the syncytium then use the “CLEAN” function of the **microinjector** to introduce the injection mix at approximately 99 psi until a clear gonadal flow of injected liquid is observed to ensure reagents accessed germ cells in the gonadal syncytium.

3.3.6 If the second gonadal arm is accessible, move the stage to position the needle to inject into the second arm.

#### 3.4. Nematode recovery post injection

3.4.1 Immediately after injection, resuspend the injected animal in 1 μL-2 μL of M9 buffer on the agarose pad.

3.4.2 Using a thin platinum wire, pick up the injected nematode off the agarose pad and wash off the oil using M9 buffer.

3.4.3 Recover the nematodes on NGM plates with a lawn of *C. aquatica* bacteria (NGM plate preparation: see Section 1.1) which increases the survival rate of the nematodes post-injection^14^. Incubate at 25°C overnight.

3.4.4 The next day, isolate each injected nematode (P0) onto an individual NGM plate seeded with *X. griffiniae*. Let P0 animals produce F1 progeny overnight (either by laying eggs or bagging) for phenotyping and genotyping (See section 4).

### 4. Screening to confirm CRISPR-Cas9 mediated knockouts

#### 4.1. Nicotine-based (CAUTION) screen of F1 for twitching phenotype when using *unc-22* as a co-CRISPR marker

4.1.1 Transfer F1 progeny from injected P0 animals into a spot of 2% Nicotine solution.

4.1.2 Pick an individual twitching animal from the nicotine spot and immediately recover the animal on NGM agar seeded with *X. griffiniae*. Isolate each twitching F1 onto individual plate and let it produce F2 animals for genotyping (See Section 4.2)

#### 4.2. Genotyping

4.2.1 Pick eight F2 or F3 progeny from each F1 twitching line for genotyping: these animals are likely to include both +/*mut* heterozygous and *mut/mut* homozygous animals.

4.2.2 Add 20 μL Proteinase K (10 mg/mL) to 180 μL worm lysis buffer (50 mM KCl, 10 mM Tris pH 8.0, 2.5 mM MgCl2, 0.45% NP-40, 0.45% Tween 20, 0.01% gelatin).

4.2.3 Add nematodes to 10 μL of the worm lysis buffer and Proteinase K mix in PCR tubes. Incubate at 65°C for 10 minutes then 95°C for 15 minutes to extract genomic DNA.

4.2.4 Genotype the nematodes by PCR-amplifying the Cas9 target region following manufacturer’s instructions, using 1-2 μL of lysate from 4.2.2. It is important to allow for ample space, if possible (at least 300 bp), between the Cas9 cut sites and the primer binding sites in case large deletions occur. Agarose gel electrophoresis can be used to confirm successful amplification before sequencing.

4.2.5 Clean up the PCR product following the manufacturer’s instructions is necessary for accurate Sanger sequencing trace analysis.

4.2.6 Sequence the PCR amplicons generated in 4.2.4 using Sanger sequencing and analyze the chromatograms using the ICE program ^29^.

4.2.7 If mutant alleles are confirmed in 4.2.6, go back to the original F1 twitching plate and isolate at least eight hermaphrodites onto individual plates. Allow each single nematode to produce progeny then mass genotype the progeny to confirm homozygosity. Homozygous lines can be maintained in the laboratory as outlined in section 5.

### 5. CRISPR lines maintenance in laboratory

There are three main methods for line maintenance. Trehalose-DMSO based simple cryopreservation protocol to back up the wildtype and mutant strains (section 5.1). For lines that are not viable after cryopreservation, use i*n vivo* (*Galleria* waxworm) growth (section 5.2).

#### 5.1. Trehalose-DMSO cryopreservation of wild-type and mutant lines

Using a trehalose-DMSO cryopreservation method ^4^, *S. hermaphroditum* can be stably stored long term at -80°C and recovered on NGM medium. Nematodes are frozen in 2 mL cryogenic vials. Optimal freezing is achieved through gradual cooling using a polystyrene foam container.

5.1.1 Grow *S. hermaphroditum* on NGM seeded with *X. griffiniae* [see section 1.1] until the majority of the population is J1 or bagging hermaphrodites. Note: avoid infective juvenile (IJ) stages since they do not freeze well using this method.

5.1.2 Wash nematode worms off from NGM agar plates in M9 buffer and collect in 15 mL conical tubes up to 5 mL mark. Concentrate by centrifugation using the floor centrifuge (1372 x g for 1 - 2:30 minutes is sufficient).

5.1.3 Remove supernatant and wash once in 5 mL room temperature Trehalose-DMSO freezing buffer by centrifugation following the parameters in 5.1.2.

5.1.4 Re-suspend worms in 5 mL room temperature Trehalose-DMSO freezing buffer. Incubate at room temperature for 30 minutes with the tube on its side. This increases the rate of survival.

5.1.5 Dispense 1 mL per tube in 4 cryogenic vials.

5.1.6 Place vials in polystyrene foam boxes in the -80°C freezer.

5.1.7 1 week later, test thaw one of the vials.

5.1.8 Transfer the rest of the vials into -80°C freezer boxes and store permanently.

**5.1.9 The test thaw or recovery *S. hermaphroditum* from frozen stocks:**

5.1.9.1 Remove a vial from the freezer and allow it to thaw completely at room temperature.

5.1.9.2 Pour the liquid contents onto a fresh NGM plate. Note that addition of symbiotic bacterium is not necessary since the frozen stock usually contains sufficient symbiotic bacteria. Worms should start recovering within minutes to hours.

5.1.9.3 Observe worm movement and reproduction. After a day or two, transfer several worms to new NGM plates seeded with *X. griffiniae*.

#### 5.2. *in vivo* maintenance through *Galleria mellonella*

For *in vivo growth*, propagate *S. hermaphroditum* mutant lines through *Galleria mellonella* (wax worm) 5^th^ instar larvae. A detailed and similar protocol of entomopathogenic nematode natural propagation through insects, and other media for EPN growth can be found in (McMullen, J.G., 2nd, Stock, S.P, 2014 ^30^). In brief:

5.2.1 Allow worms to starve and become IJs on NGM plates.

5.2.2 Use M9 buffer to remove IJs from the NGM plates.

5.2.3 Transfer IJ to infect *Galleria* insects using the white trap.

5.3 *in vitro* maintenance on NGM plates

5.3.1 Seed NGM plates with approximately 100 μL of fresh liquid culture of the *S. hermaphroditum* symbiont, *X. griffiniae*.

5.3.2 Allow the bacterial patch to fully dry before transferring the nematodes onto the plate.

5.3.3 Store the plates at 25 °C and transfer the nematodes to freshly seeded plates approximately once a month.

## REPRESENTATIVE RESULTS

Two CRISPR RNAs were designed (identified by the Pam site number – Pam 3 and Pam 5) targeting the *S. hermaphroditum* homologue of *unc-22* at previously published Pam sites [Figure 2A, ^14^]. Each crRNA complexed with tracrRNA to form a single guide RNA (sgRNA) and was incorporated into Cas9 to form the ribonucleoprotein as described in section 3.1. In this protocol, two gRNAs were designed to target the same gene, however, if using the co-CRISPR method developed for use in *C. elegans* ^31^, crRNA #2 can be replaced by *unc-22* crRNA. Alternatively, the *unc-22* crRNA can be added to the two targeting the gene of interest making sure to preserve the 1:1 ratio between total crRNA and tracrRNA.

**Figure 2:**
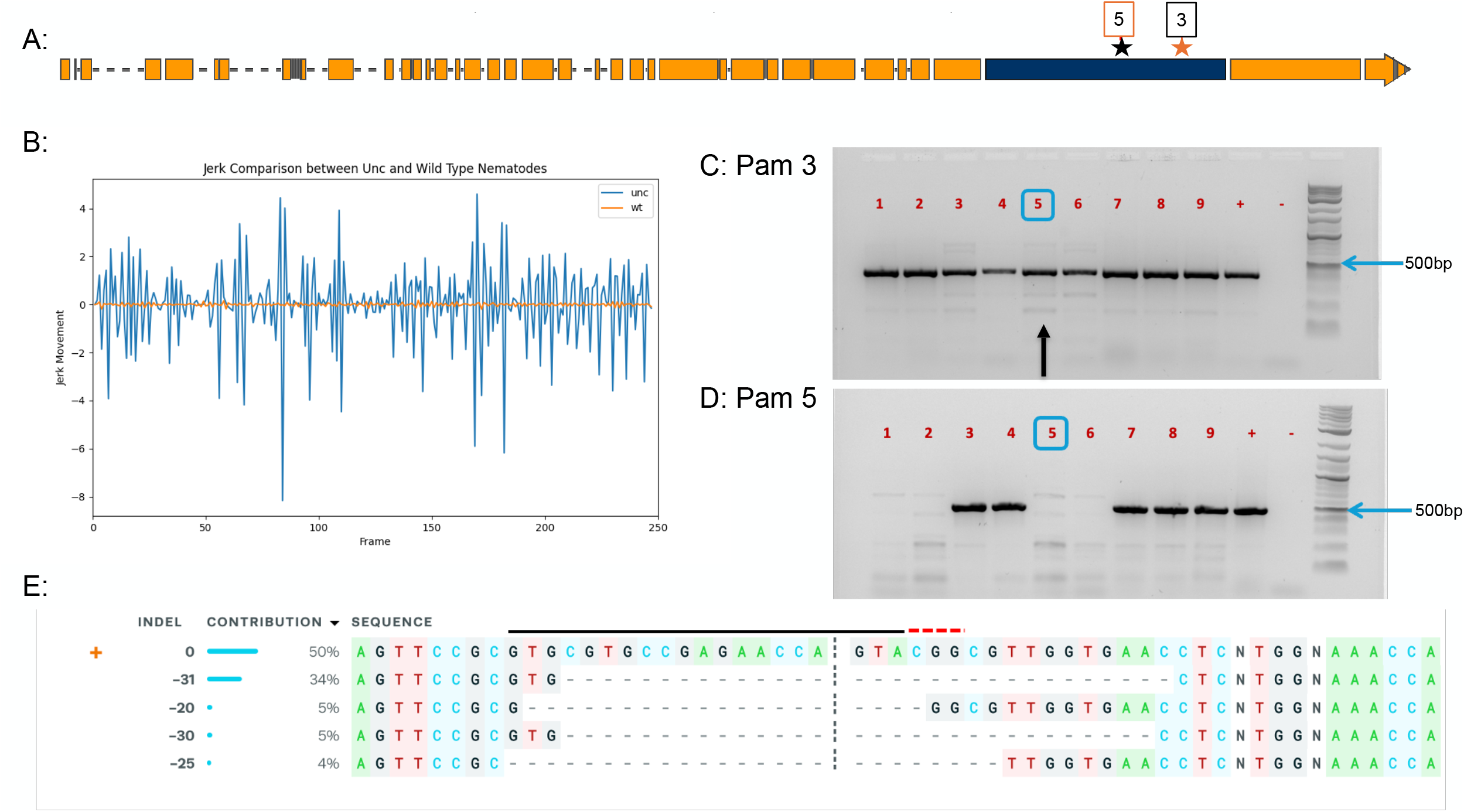
Representative results of CRISPR-Cas9 genome editing in *S. hermaphroditum*. (A): The gene structure of *S. hermaphroditum unc-22* homologue. Blue box: the targeted exon. Black star: Pam 3 target site. Red star: Pam 5 target site. (B): Phenotypic analysis of a representative twitching progeny based on jerk movement when treated with 2% nicotine. Blue: a twitching adult later confirmed to be *unc-22* mutant. Orange: a wild-type adult paralyzed by 2% nicotine. (C and D): Genotyping of nine independent F3 twitching lines (1-9) from two injected P0 animals. Each sample contains four F3 progeny from the same F1 line. Primers were used to amplify the genomic region around Pam 3 (C) and Pam 5 (D). ‘+’: wild-type control; “-”: a negative control with no DNA loaded. Blue boxes: example of a confirmed heterozygous sample amplicon on an agarose gel. Black arrow in panel (C): the amplicon analyzed in panel (E). (E): ICE analysis of *unc-22* mutant at Pam 3. PCR sample 5 from panel (C) (black arrow) was sequenced, and analyzed using ICE, showing 50% of the alleles being wild-type, while various deletion mutations (ranging from 20 bp-31 bp) were detected in the genomic DNA.

The day prior to microinjections, well-fed early J3 nematodes with visible gonad arms (Fig. 1A) were picked onto a fresh plate seeded with the symbiotic bacterium *Xenorhabdus griffiniae* and allowed to grow into young adults overnight at room temperature (Fig. 1B). On the day of injections, young adults (Fig. 1B) carrying few to no fertilized embryos were isolated onto a new petri-dish for injection. Based on observation, younger nematodes had a relatively more penetrable cuticle, however, were less likely to recover after injection. The needle was made [see section 2.3], loaded with the injection mix and broken open as described in section 3.2. The nematode was washed in M9 buffer and immobilized in a droplet of halocarbon oil on the agarose pad [section 2.2]. At the injection site [Fig. 1C], external pressure from the microinjector was used to create a visible liquid flow in the gonad syncytia, ensuring the germline nuclei were bathed in the injection mix [section 3.3]. Where possible, both gonads were injected to increase the probability of obtaining edited F1 progeny.

The injected nematode (P0) was quickly recovered by washing in M9 buffer and incubated overnight on NGM plates seeded with *Comamonas aquaticus* bacteria for overnight recovery [section 3.4]. Approximately 40%–90% of injected P0 animals recovered soon after injection, and 20%–90% of injected P0 nematodes survived overnight on *C. aquatica*. [Table 1]. Individual P0 animals that survived were then isolated onto NGM petri-dishes seeded with *X. griffiniae* with a transfer rate of 20%–90%. Nicotine screening for the *unc-22* deletion phenotype was performed on F1, F2 and F3 animals to determine the heritability of the deletion [section 4.1]. *S. hermaphroditum* nematodes with the *unc-22* gene deletion twitch when exposed to 2% nicotine, while wildtype nematodes become paralyzed which allows for screening and selection of animals with putative deletions ^28^. In this experiment, up to 64% of F1 nematodes isolated from injected P0 produce consistently twitching F2 progeny (Table 1). For the ease of screening co-CRISPR injections, individual twitching nematodes can be monitored and quantified using imaging and tracking systems, with one example shown in (Fig. 2B).

To confirm the genotype of twitching progeny, genomic DNA was extracted from F3 nematodes to genotype the *unc-22* locus [section 4.2]. By PCR-amplifying the genomic region surrounding targeted sites, multiple faint bands were observed indicating small and large insertions and deletions (indels) (Fig. 2C and Fig. 2D), some of which were confirmed by sequencing analysis (Fig. 2E). These data suggest CRISPR-Cas9 created double-stranded breaks at the genomic target sites which were repaired *via* the non-homologous end-joining (NHEJ). To confirm the specific modification of genome at the nucleotide level, PCR-amplified products from the target sites were analyzed using Sanger sequencing: alignment of the chromatograms from both the twitching and wildtype nematodes and allowed for identification of insertions or deletions at the target sites (Fig. 2E). Once confirmed by phenotyping and genotyping, the CRISPR mutant lines can be maintained in the laboratory by passing through the insect *Galleria mellonella* or preserved by cryopreservation [section 5]. If using *unc-22* as a co-CRISPR marker, see Fig. 3 for an illustration of a suggested strategy to isolate F2 and F3 progeny carrying mutations in the target gene.

**Figure 3:**
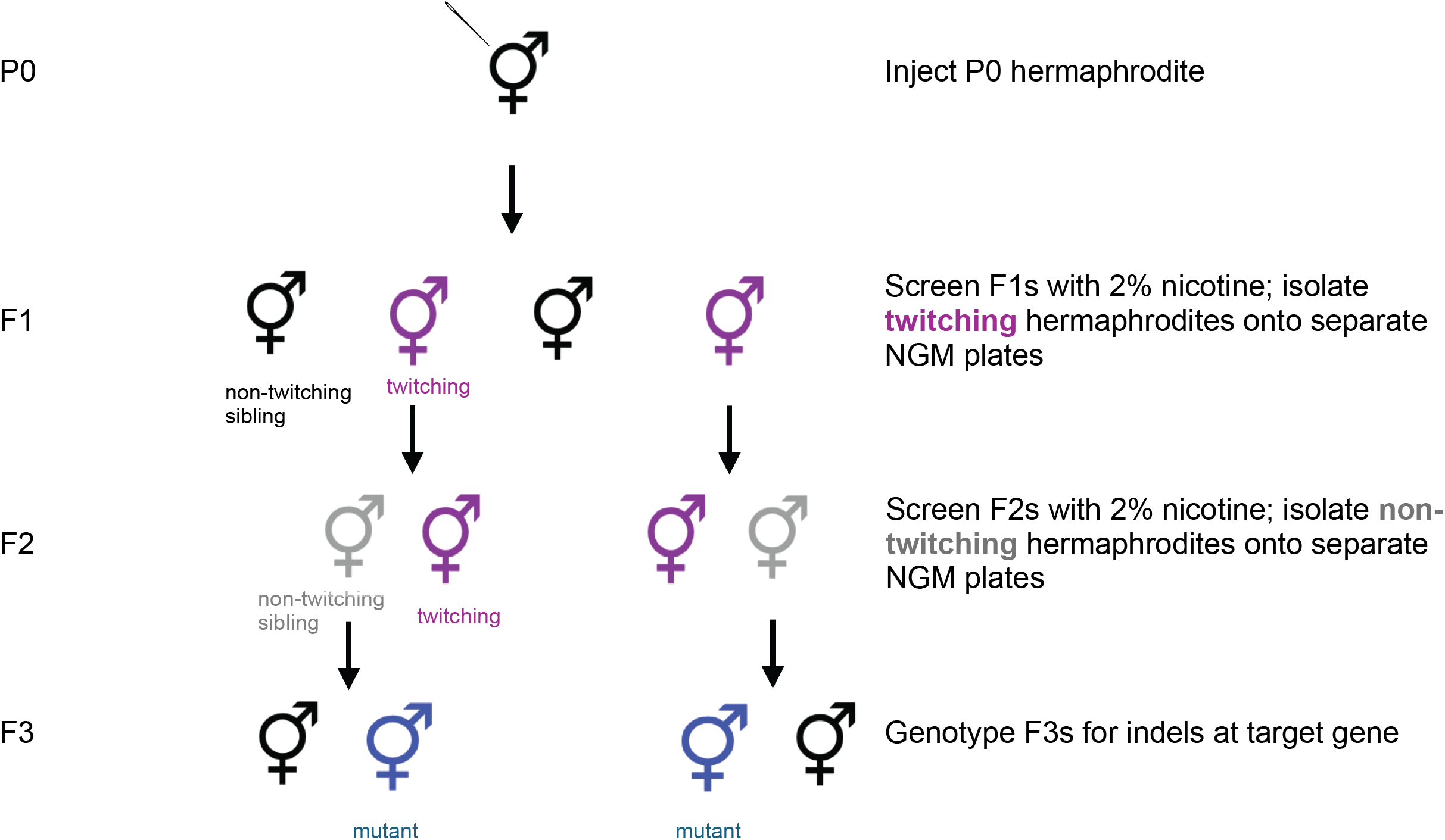
Co-CRISPR screening workflow to isolate mutants in the gene of interest. When using *unc-22* as a co-CRISPR marker, screen the F1 nematodes with 2% nicotine and isolate the twitching F1s onto separate NGM plates seeded with the symbiont *X. griffiniae*. Screen the F2 progeny of twitching F1s and isolate non-twitching F2s onto separate plates. Allow the F2s to produce progeny, then genotype F3s to determine if there is a knockout of the targeted gene. Note that the non-twitching siblings of twitchers could potentially have indels at the gene of interest, and these can also be screened by genotyping.

## FIGURE AND TABLE LEGENDS

**Table 1**: Representative results of the *S. hermaphroditum unc-22* homologue modification via microinjection and rate of success as evidenced by twitching progeny.

*Survival was determined by movement of the pharyngeal pump approximately 30 minutes after microinjection.

## DISCUSSION

Traditionally non-tractable animals offer valuable opportunities to study eukaryotic gene functions in the relevant context of the animal’s ecological niches, such as in the presence of their parasitic host or natural microbiome. Alternative methods of genetic manipulation in EPNs have largely been RNAi-based^13, 32^. While this technology allows for the study of gene function, it is limited by the transient nature of RNAi as it is not reliably heritable, as well as variable levels of gene silencing underlining the need for a more stable form of genetic manipulation^33, 34^. This is the first protocol detailing precise genome editing in Steinernematids, one of the two families of EPNs in the order Rhabditida, as demonstrated in (Cao M 2023^14^).

Handling of the nematodes is crucial for successful CRISPR-Cas9 mediated gene knockouts using this protocol. If the nematodes are not well fed, the gonads collapse and are difficult to penetrate with the needle. The injection volume should be adjusted by breaking open the needle further or adjusting injection pressure to ensure the gonad is flushed with the injection mix during microinjection. Prompt recovery of the injected nematode from the agarose pad to the *C. aquatica* recovery plate is critical for survival after injection, and this speed improves with practice. After microinjection, care should be taken not to expose the wounded nematode to its symbiont as post-injection recovery on *X. griffiniae* is low^14^.

Clogging of the needle during microinjection is a common problem encountered which can sometimes be resolved by dragging the needle across double-sided tape submerged in halocarbon oil. Rinsing the nematodes well in M9 before injection prevents the symbiont bacteria from clogging the needle. Alternatively, breaking open the needle further may resolve the blockage, at the risk of breaking it open too far resulting in free flow of the injection mix. The switch from borosilicate^14^ to quartz capillaries for making the microinjection needles makes it less challenging to inject slightly older adults which tend to have a thicker cuticle, hence improving survival.

A significant limitation of this protocol is that CRISPR-based transgene insertion is currently challenging and requires further optimization. Homology-directed repair (HDR) of double-stranded DNA breaks caused by Cas9 has been difficult to achieve in *S. hermaphroditum* using short ssDNA^14, 35^. Precise transgene insertion is an important technique in the genetic toolkit of an organism that allows for genome modification with high fidelity such as single amino acid modifications or reporters for gene expression^36^. Therefore, the development of a consistent CRISPR-Cas9 mediated knock-in protocol would be of great benefit to the scientific communities of parasitology, microbial symbiosis and beyond.

## Supporting information

Materials list

## ACKNOWLEDGMENTS

We thank Margaret McFall-Ngai and Edward Ruby for use of their confocal microscope. We thank Alexis (Cody) Hargadon and Grischa Chen for training on confocal microscopy, and Carly R. Myers for help with obtaining the pictures and analysis of twitching videos. We also thank Stephanie Hampton for allocating funds for purchase of the microinjection system. Carnegie Institution for Science supported this work.

